# Identification by MALDI-TOF MS of Coleopteran insect pests collected in the field

**DOI:** 10.1101/2025.08.07.669236

**Authors:** Dikra Hamadouche, Fabien Fohrer, Adama Zan Diarra, Jean-Michel Bérenger, Philippe Parola, Lionel Almeras

## Abstract

In the context of museum and cultural heritage protection, rapid and accurate identification of Coleopteran pests is crucial. This study evaluated the effectiveness of MALDI-TOF MS in identifying seven species of Coleoptera collected between 2014 and 2023 from 17 French cultural sites. A total of 273 specimens (205 field-collected and 68 laboratory-reared) were analysed. Although reproducible and species-specific MS spectra were obtained, intra-species reproducibility was higher for laboratory reared specimens frozenly-stored compared to counterparts field collected and stored at room temperature. Two successive blind tests were done against the reference MS database including, firstly (ie, database 1, DB1), only laboratory-reared Coleoptera MS spectra, and secondly, those of DB1 upgraded with MS spectra from field collected specimens (ie, BD2). Correct identification at the species were obtained for MS spectra query against DB1 for species which possess homolog in the database. However, the spectra reaching threshold for relevant identification were coming from essentially laboratory-reared Coleoptera. An upgrading of the reference MS database (ie, BD2), improved identification performance for field specimens, although limitations remained for species with MS spectra of low peak diversity like *Pentarthrum huttoni*. In parallel, three specimens per species and storing mode were submitted to molecular identification using COI and 16S markers. For field samples, the rate of successful PCR product sequencing was lower than 10%, which was likely attributed to DNA degradation of these samples stored long term at room temperature. Overall, the study confirmed that MALDI-TOF MS appeared as a promising tool for Coleopteran pest identification, especially when reference spectra matched both species and preservation mode.

## 1. Introduction

The conservation of cultural heritage is a major challenge for museums, libraries, archives, and other heritage institutions (1). Among the many threats to these valuable assets, infestation by wood-boring and keratin-feeding insects represents a significant danger (2). These pests can cause irreversible damage to wooden objects, textiles, ancient manuscripts and book bindings, thereby compromising their long-term preservation (3).

One of the most pressing challenges in heritage conservation is the damage caused by certain insect species, which play a particularly destructive role in the degradation of organic materials found in historical collections (4). Among them, *Stegobium paniceum* and *Anobium punctatum* pose a significant threat to libraries in Central Europe. These pests burrow into books, leaving distinctive holes in the pages, and can severely damage book bindings and ancient manuscripts (5). *S. paniceum,* also known as the bread beetle or the drugstore beetle, reproduces rapidly, accelerating infestations and increasing destruction (6). Meanwhile, *A. punctatum*, commonly known as the furniture beetle, primarily targets wooden structures, weakening beams and antique furniture through its tunneling activity, further compromising the integrity of cultural heritage (7). In addition to these beetles, *Mezium affine*, *Oligomerus spp*., and *Anthrenus verbasci* (the varied carpet beetle) also feed on organic materials and can cause significant structural damage to artworks, antique furniture, and historical documents.

Accurate identification of these pests is crucial for implementing effective conservation strategies (8). Traditionally, identification has been based on morphological characteristics, a method that requires extensive entomological expertise (9). However, this approach has several limitations, as it is often time-consuming (10) and hindered by the morphological similarities between species (11). Moreover, identifying larvae—often the primary agents of degradation— remains particularly challenging, as most available identification keys primarily focus on adult insects (12). In light of the limitations of morphological identification previously mentioned, molecular identification techniques provide a precise and reliable alternative to address the challenges associated with analysing hard-to-characterize specimens (13). Among these techniques, DNA barcoding (14) and metabarcoding technologies, also known as high-throughput sequencing (HTS), offer significant advantages for species identification (15). DNA barcoding relies on sequencing a specific gene, typically mitochondrial cytochrome c oxidase I (CO1), to differentiate species from biological samples (16). When mitochondrial DNA sequences are unavailable, alternative genetic markers, such as 16S rDNA, are often used as reliable substitutes (17). However, a major limitation of this technique is the frequent occurrence of taxonomic errors in published DNA sequence databases (18–20). On the other hand, metabarcoding is a method that enables the simultaneous identification of multiple arthropod species from a single pooled sample containing a mixture of individuals (21,22). This approach has been used to analyze the contents of entomological traps and characterize arthropod communities (23). This approach is both rapid and cost-effective, making it increasingly relevant in the field of ecology (24). However, despite its clear advantages, metabarcoding remains a relatively recent method, with several challenges yet to be addressed (25). The reliability of species identification through genetic sequences depends on the quality and availability of reference databases. Some databases are precise and complete, while others are incomplete or less reliable (26). As a result, metabarcoding analyses can be prone to false negatives, where species go undetected despite being present, due to insufficient sampling or biological and technical limitations that lead to detection errors (27). Therefore, developing alternative approaches that tackle these challenges and offer more accessible solutions is crucial for advancing routine species identification. To mitigate the limitations of both morphological and molecular identification methods, the development of new strategies has become essential.

A recent study demonstrated the effectiveness of matrix-assisted laser desorption/ionization time-of-flight mass spectrometry (MALDI-TOF MS) for identifying adults, larvae, and exuviae from eight Coleopteran species (28). Reproducible intra-species MS spectra were obtained for each developmental stage, confirming the specificity of spectral signatures and supporting the use of this technique for specimen classification. The spectral profiles for each stage were added to a home-made arthropod MS reference database. This study represents the first demonstration of the successful application of MALDI-TOF MS as a reliable, rapid, and cost-effective tool for identifying different developmental stages of Coleopteran pests affecting France’s cultural heritage. However, the specimens used in this pioneering study were all reared in laboratory conditions, which may differ significantly from those encountered in the natural environment (28).

The aim of this study was to evaluate the application of MALDI-TOF MS for monitoring Coleopteran samples collected in the field essentially in museums. To this end, seven Coleopteran species were tested, among which uniquely one species was common with our previous study (28). The samples analysed consisted uniquely of Coleopterans at adult stage collected in the field over the last 1 to 10 years and stored at room temperature. The specimens came from 17 collection sites in France. The study was designed to assess the effect of preservation duration and mode. Moreover, the consequences of the geographical origin on the MS profiles obtained among specimens from the same species were examined. An additional goal was to generate reference MS spectra for newly identified beetle species, in order to improve the database for future species identification.

## 2. Materials and methods

### 2.1. Specimen collection, selection and morphological identification

All Coleopterans included in the present study, were collected in the field this last decade in France by members of the “Centre Interdisciplinaire de Conservation et de Restauration du Patrimoine” (CICRP). Among the Coleopteran specimens available in the CICRP collections, those included in this study were selected according to the following criteria: (1) accurate morphological identification at the species level; (2) preference for the adult stage; (3) origin from cultural heritage sites; (4) inclusion of species not represented in previous studies; (5) storage duration not exceeding 10 years; and (6) consistent storage conditions across all specimens.Then, seven different species collected from 17 sampling sites, primarily in museums, and other collection areas were included in the present work (Table 1). Among the seven species, two species were either collected in the field or freshly sampled coming from rearing done as previously described (28). The fresh specimens were used as Coleopteran positive controls of the different experiments performed. The field samples were gathered over a period spanning from 2014 to 2023 using manual collection. All these specimens were stored at ambient temperature until further analysis.

**Table 1.**
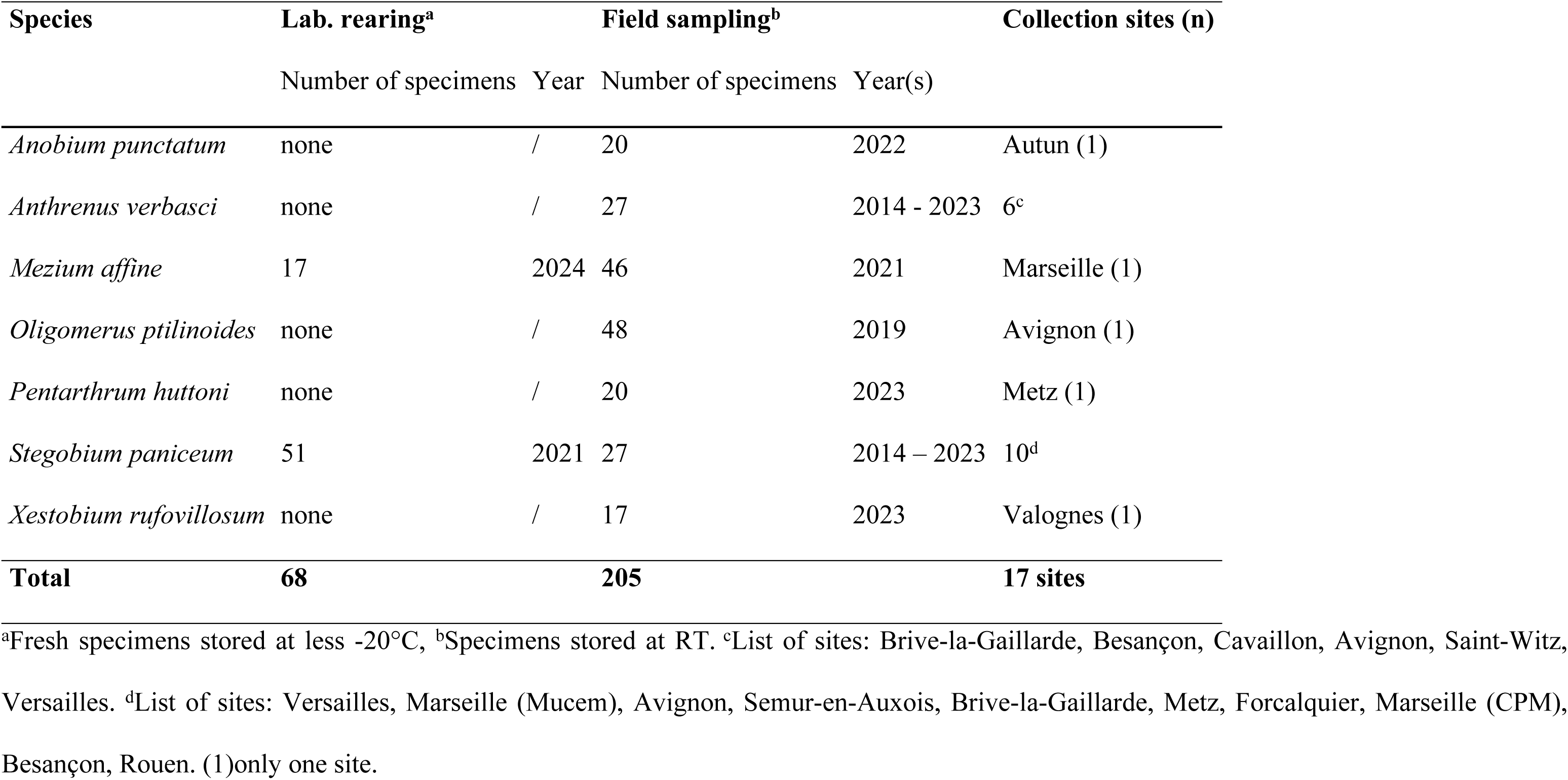
Details about the Coleopterans selected from laboratory and field origins.

### 2.2. Dissection and sample preparation

Prior dissection, each sample was washed for two minutes with 70% ethanol, followed by two successive baths of distilled water before to be dried on sterile filter paper. The samples were then dissected under a Leica ES2 10x/30x stereomicroscope using sterile slide, scalpel and pliers for each specimen. Based on a previous study (28), for adult Coleopterans, the thorax and legs were the body part selected for MS analysis which requires a standardization of the methodology to ensure consistent results. In order to minimise variations in MS spectra within species associated with their feeding habits (29,30), the abdomens of adult specimens were systematically excluded from the MS analysis and used for molecular biology studies. For MS analysis, thorax and legs from each specimen was placed individually in 1.5 mL tubes and crushed for 3 minutes with a Tissue-Lyser (Qiagen, Germany) at a frequency of 30 hertz, using a pinch of glass beads (1.0 mm) (Sigma Aldrich, St. Louis, Missouri, USA) in 40 µL of a buffer mixture composed of 50% (v/v) 70% formic acid (Sigma-Aldrich, Lyon, France) and 50% (v/v) acetonitrile (Fluka, Buchs, Switzerland). This homogenization step was repeated two times for a total of three cycles, as described previously (31). Following homogenization, a brief centrifugation was performed for one minute at 2000x g. The samples were immediately loaded onto MS plate.

### 2.3. Sample loading on the target plate and MALDI-TOF MS settings

For each sample, one μL of the supernatant was spotted in quadruplicate onto a steel target plate (Bruker Daltonics, Wissembourg, France). After spots drying at room temperature, each spot was overlaid with 1 μL of a matrix buffer, composed of α-cyano-4-hydroxycinnamic acid CHCA at 10 mg/mL (Sigma, Lyon, France), 50% acetonitrile (v/v), 2.5% trifluoroacetic acid (v/v) (Aldrich, Dorset, UK), and HPLC-grade water (32). The matrix buffer was air-dried at room temperature before the target plate was placed into the mass spectrometer for analysis. Legs and thorax from laboratory reared *Aedes albopictus* mosquitoes, prepared as previously described (33), were used loaded on each plate as a quality control of sample preparation and MS spectra acquisition. Moreover, to control the absence of impurities in the matrix buffer, matrix buffer alone was loaded in duplicate onto each MALDI-TOF plate (i.e., negative control).

### 2.4. MALDI-TOF MS parameters

Protein mass profiles were obtained using a Microflex LT MALDI-TOF Mass Spectrometer (Bruker Daltonics, Germany), operating in linear positive-ion mode at a laser frequency of 50 Hz, with a mass range of 2-20 kDa. The accelerating voltage was set to 20 kV, and the extraction delay time was 200ns (34). Each spectrum was generated from ions produced by 240 laser shots across six regions of the same spot, automatically acquired using the AutoXecute method in the flexControl v2.4 software (Bruker Daltonics).

### 2.5. Analysis of spectra

The quality of the MS spectra obtained from each specimen was initially assessed through visual inspection using ClinProTools v2.2 and Flex analysis v.3.3 software (Bruker Daltonics) (35). Intra-species reproducibility and inter-species specificity were examined using cluster analysis (MSP dendrogram) using MALDI-Biotyper v3.0 software (Bruker Daltonics). To estimate MS spectra distance between species and/or storing mode per species, coefficient correlation index (CCI) matrix was calculated using MALDI-Biotyper v3.0. software with default settings (mass range 3.0-12.0 kDa; resolution 4; 8 intervals; auto-correction off). Higher correlation values (expressed by mean ± standard deviation – SD) reflecting higher reproducibility for the MS spectra.

### 2.6. Creation of the reference databases and blind tests

Two databases were created for this study to assess the impact of storing mode of specimens on the accuracy of species identification. The DB1 was composed of reference MS spectra from several arthropod species (PMID: 35623553), including thorax and legs MS spectra uniquely from the reared Coleopteran species freshly submitted to MALDI-TOF MS or stored at less - 20°C. Then, DB1 included all reference MS spectra from the previous study (28) which comprised mass profiles of six Coleopterans species at adult stage (ie, *Lasioderma serricorne, Attagenus smirnovi, Trogoderma versicolor, Reesa vespulae, A. verbasci, Anthrenus pimpinellae*), more two new species (ie, *M. affine, S. paniceum*) from the present study. The DB2 contained all MS spectra from reared Coleopterans (ie, DB1), more those from field collection (ie*, M. affine, S. paniceum, A. punctatum, Oligomerus ptilinoides, P. huttoni, Xestobium rufovillosum*) (Table 2). MS spectra from three specimens per species were incorporated into our in-house reference database, which includes over two thousand spectra from various arthropod species (36). MS spectra included in the database were generated through an unbiased algorithm that analyses peak position, intensity, and frequency data using MALDI-Biotyper v3.0 software (Bruker Daltonics) (37). The effectiveness of MS profiling for Coleopteran species identification was evaluated by conducting blind tests on the remaining spectra, by comparing them to the MS reference database (ie, DB1 or DB2). The accuracy of species identification was determined using log score values (LSVs), which ranged from 0 to 3, and it was calculated using a biostatistical algorithm from the MALDI Biotyper software v.3.0. (Bruker Daltonics). According to previous studies (33,38), a LSV upper than 1.8 should be reached to consider species identification as reliable. Data were analysed with Prism software v.7.00 (GraphPad, San Diego, CA, USA).

**Table 2.** Identification of Coleopteran species using MALDI-TOF MS according to reference spectra DB contains.

| Morphological ID | Sampling origin | Number of samples | DB1<sup>a</sup> | Blind tests against DB1 | Concordance of MS identification with morphological ID | Range of LSVs | of LSVs ≥ 1.8 N (%)<sup>c</sup> | DB2<sup>b</sup> | Blind tests against DB2 | Concordance of MS identification with morphological ID | Range of LSVs | of LSVs ≥ 1.8 N (%)<sup>c</sup> | Paired comparisons of LSVs (W-test)<sup>e</sup> |
| --- | --- | --- | --- | --- | --- | --- | --- | --- | --- | --- | --- | --- | --- |
| *Ano. punctatum* | Field | 20 |  | 20 | 0 | 0.62 - 1.37 | 0 | 3 | 17 | 17 (100%) | 1.26 - 2.31 | 8 (47.1%) | *p*<0.0001 |
| *Ant. verbasci* | Field | 27 |  | 27 | 27 (100%) | 0.78 - 1.89 | 1 (3.7%) | 3 | 24 | 24 (100%) | 1.02 - 2.13 | 8 (33.3%) | *p*<0.0001 |
| *M. affine* | Bread | 17 | 3 | 14 | 14 (100%) | 1.65 - 2.30 | 13 (92.9%) | 3* | 14 | 14 (100%) | 1.65 - 2.30 | 13 (92.9%) | *n.d.* |
|  | Field | 46 |  | 46 | 41 (89.1%) | 0.64 - 1.63 | 0 | 3 | 43 | 43 (100%) | 0.83 - 1.89 | 3 (7.0%) | *p*<0.0001 |
| *O. ptilinoides* | Field | 48 |  | 48 | 0 | 0.70 - 1.34 | 0 | 3 | 45 | 45 (100%) | 1.13 - 2.13 | 36 (80.0%) | *p*<0.0001 |
| *P. huttoni* | Field | 20 |  | 20 | 0 | 0.50 - 1.11 | 0 | 3 | 17 | 15 (88.2%) | 0.74 - 1.57 | 0 (0.0%) | *p*<0.001 |
| *S. paniceum* | Bread | 51 | 3 | 48 | 48 (100%) | 1.88 - 2.71 | 48 (100%) | 3* | 48 | 48 (100%) | 1.88 - 2.71 | 48 (100%) | *n.d.* |
|  | Field | 27 |  | 27 | 25 (92.6%) | 0.56 - 1.75 | 0 | 3 | 24 | 24 (100%) | 0.66 - 1.80 | 1 (4.2%) | *p*<0.0001 |
| *X. rufovillosum* | Field | 17 |  | 17 | 0 | 0.42 - 1.02 | 0 | 3 | 14 | 14 (100%) | 1.03 - 2.11 | 4 (28.6%) | *p*<0.001 |
| Total (N)<sup>d</sup> |  | 273 (205) | 6 (0) | 267 (205) | 155 (93) |  | 62 (1) | 27 (21) | 246 (184) | 244 (182) |  | 121 (60) |  |

<sup>a</sup>DB1 includes MS spectra from other arthropods (PMID: 27070074), plus six Coleopteran species including *A. verbasci* (DOI : [10.1016/j.ibiod.2024.106033](https://doi.org/10.1016/j.ibiod.2024.106033)), more those from laboratory-rearing listed in the table. <sup>b</sup>DB2 includes MS spectra from DB1 and those from field collection listed in the table. <sup>c</sup>Proportion among concordant ID. <sup>d</sup>Number from field origin. <sup>e</sup>MS spectra already included in the DB1. <sup>e</sup>Wilcoxon matched-pairs signed rank test comparing LSVs per species against BT1 and BT2. DB, database; ID, identification; LSV, log score value; MS, mass spectrometry; n.d., not determined; N, number. *Ano.*, *Anobium*; *Ant.*, *Anthrenus*; *M.*, *Mezium*; *O.*, *Oligomerus*; *P.*, *Pentarthrum*; *S.*, *Stegobium*; *X.*, *Xestobium*.

### 2.7. DNA extraction and molecular identification

The three specimens per species, included into the MS reference database, were subjected to molecular identification. The abdomen of each specimen was incubated overnight at 56°C in 200 μL of G2 lysis buffer (Qiagen, Hilden, Germany) with 20 μL of proteinase K (Qiagen, Hilden, Germany). DNA extraction was then performed using the EZ1 BioRobot automated system (Qiagen, Hilden, Germany). The extracted DNA samples were amplified using standard PCR in an automated thermal cycler (Applied Biosystems, 2720, Foster City, USA), as previously described (39). Partial sequencing was performed for two genes, the *Cytochrome c oxidase subunit I* (COI) and the 16S ribosomal RNA gene. A 710 base-pair fragment of the COI gene was amplified using Folmer’s universal barcoding primers (LCO1490, HCO2198) (40), while the 16S rRNA gene was targeted using primers LRJ12961 and LRN13398, amplifying a 500 base-pair region (41). The PCR conditions were applied as previously described (40,42). A PCR mix without DNA was used as a negative control, while DNA extracted from laboratory-reared *Aedes albopictus* mosquitoes served as a positive control. The amplified products were then separated by electrophoresis on a 1.5% agarose gel stained with SYBR Safe™ and visualized using the ChemiDoc™ MP UV imager (Bio-Rad, Marnes-la-Coquette, France). The amplified products were visualised on 1.5% agarose gel stained with SYBR Safe then purified positive samples using a Macherey Nagel (NucleoFast 96 PCR, Düren, Germany) plate. Following purification of the positive samples, PCR products were sequenced using BigDye Terminator v1.1 and v3.1 with 5× Sequencing Buffer (Applied Biosystems, Warrington, UK). Sequencing was carried out via the Sanger method on an Applied Biosystems® 3500 Series Genetic Analyzer. The resulting chromatograms were assembled and edited using Chromas Pro 1.77 (Technelysium Pty. Ltd, Tewantin, Australia). The finalized sequences were then compared to those in the GenBank database using the BLAST algorithm (43).

### 2.8. Statistical analyses

Statistical analyses were conducted using GraphPad Prism software 7.0.0 (GraphPad Software, San Diego, CA, USA). After verifying that the LSVs in each group did not assume a Gaussian distribution, nonparametric tests were applied. A comparison of LSVs per species and storing mode between DB1 and DB2 was carried out using Wilcoxon matched-pairs signed rank test. Comparison of LSVs according to the duration of storing and collection location was done using Kruskal–Wallis tests when appropriate. All differences were considered significant at *p* < *0.05*.

## 3. Results

### 3.1. Morphological identification

A total of seven distinct Coleopteran species, collected between 2014 and 2023 from museums and cultural institutions across 17 sampling sites in France, and stored at room temperature, were selected for this study (Table 1). Among them, *A. verbasci* had already been included in a previous study demonstrating the efficiency of MALDI-TOF MS for their identification (28). The six other field collected species, including *A. punctatum, M. affine, O. ptilinoides, P. huttoni, S. paniceum and X. rufovillosum*, were studied for the first time by MALDI-TOF MS. Their morphological identification was performed by an expert in heritage pest Coleoptera (Figure 1). The opportunity to have freshly reared and field-collected specimens for two species, *M. affine* and *S. paniceum*, open the way to examine the effect of storing mode onto the resulting MS profiles. Here, a total of 273 specimens were analysed, including 68 laboratory-reared and 205 field-collected. Interestingly, field-collected specimens were particularly dry and breakable, which did not hinder their morphological identification but made dissection more challenging.

**Figure 1.**
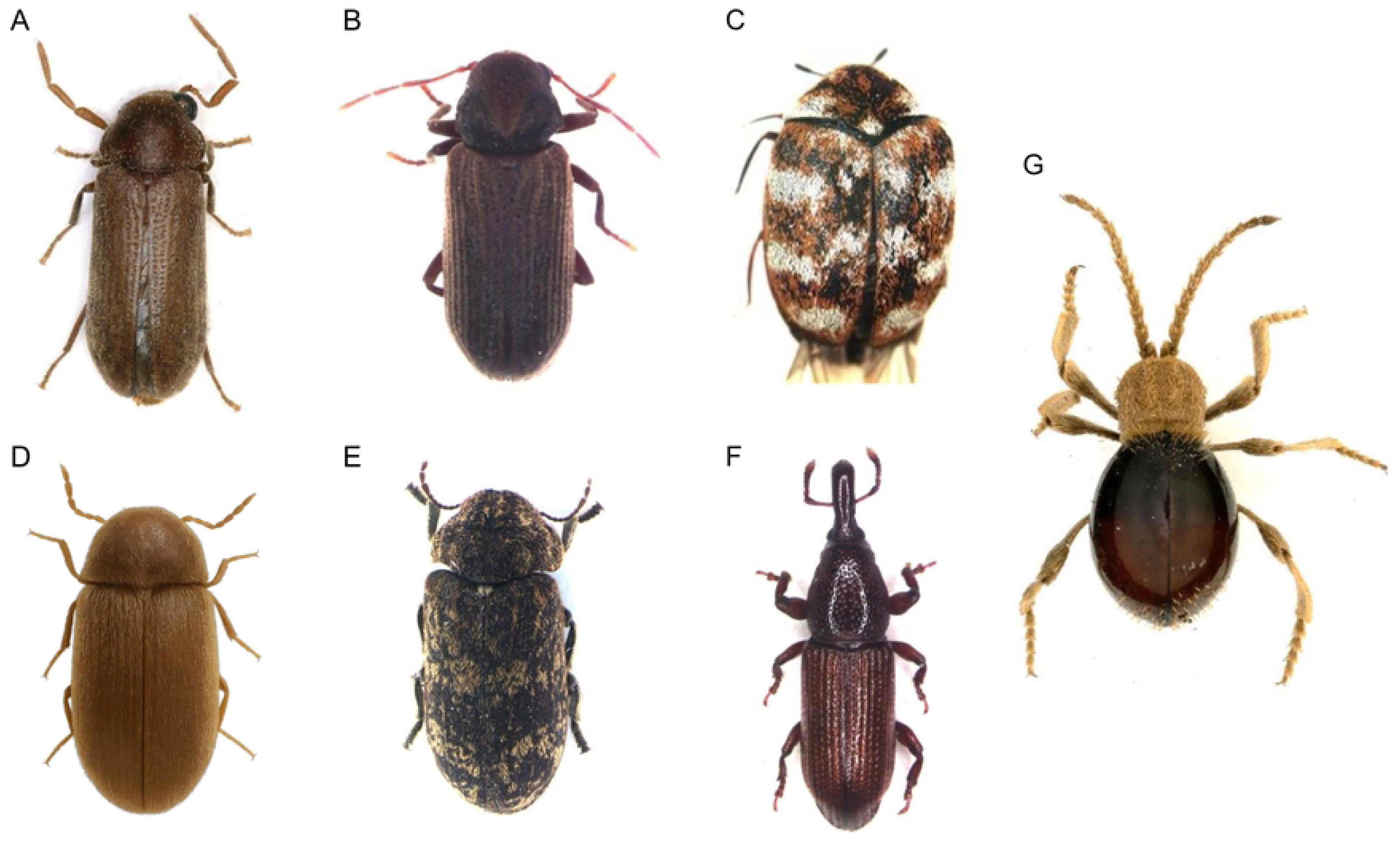
Photographs of Coleopterans species at adult stage. The seven Coleopteran species at adult stages were presented as follow: **(A)** *O. ptilinoides*, **(B)** *A. punctatum*, **(C)** *A. verbasci*, **(D)** *S. paniceum*, **(E)** *X. rufovillosum*, **(F)** *P. huttoni*, **(G**) *M. affine.* Specimens were photographed with Keyence VHX-5000 at magnification x30.

### 3.2. Specificity and Reproducibility of MALDI-TOF MS Spectra

The thorax and legs of the 273 Coleopteran specimens were subjected to MALDI-TOF MS analysis. The higher spectra intensity were obtained for both laboratory-reared species, *M. affine* and *S. paniceum*, frozenly stored. The comparison of the MS spectra from these last two species between frozen and field collected specimens stored at room temperature showed visual changes of spectral profiles according to storing mode (Figure 2A). Moreover, the MS spectra from field-collected specimens appeared generally less intensive, with greater intra-species variability (Figure 2B). This suggests that long-term storage under non-controlled conditions may impact the consistency of protein profiles used in MALDI-TOF MS analysis.

**Figure 2.**
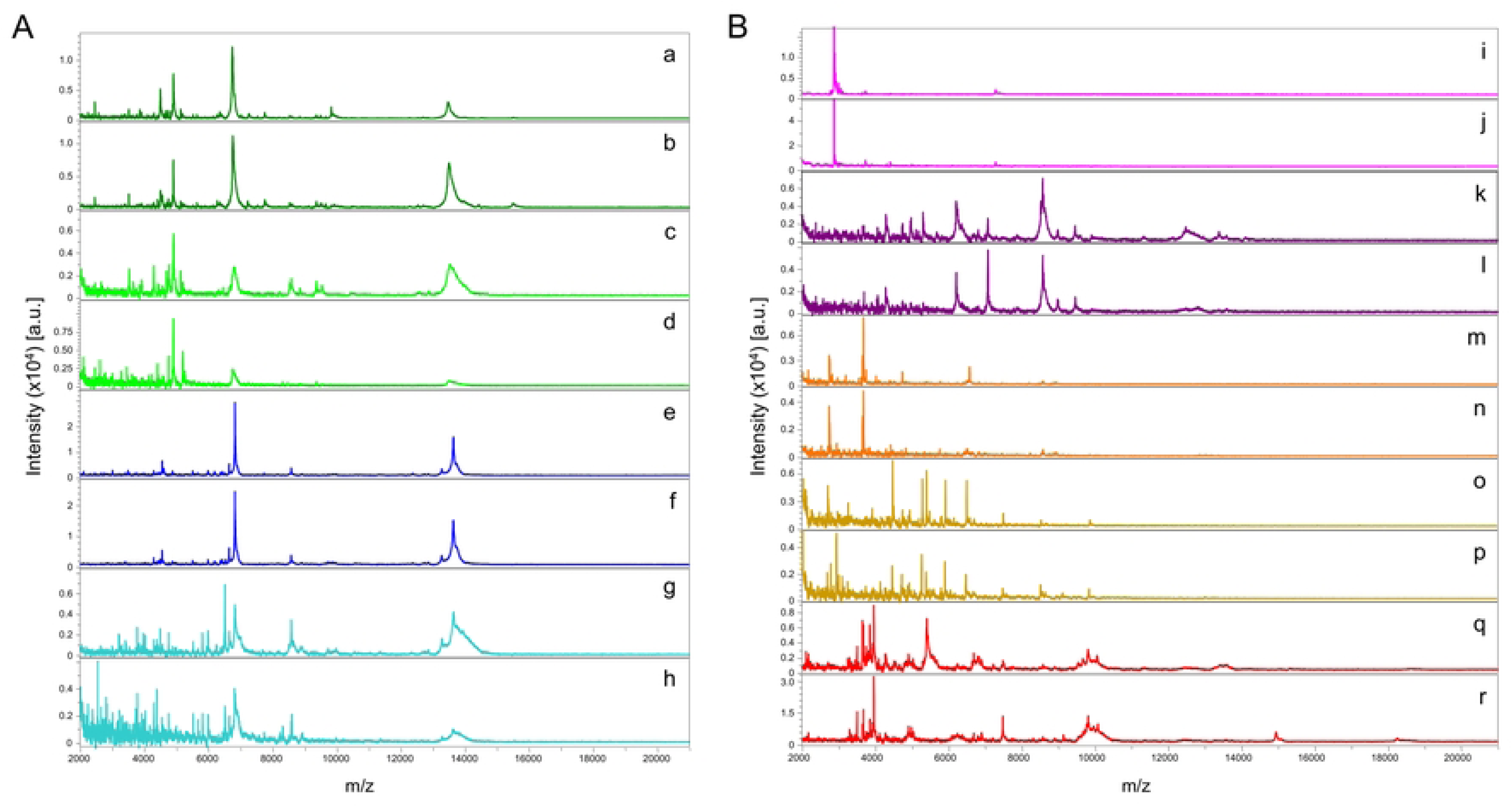
Representative MS spectra from different species of Coleoptera using flexAnalysis v.3.3. (A). Representative thorax with legs MS spectra from *M. affine* specimens laboratory-reared and stored at -20°C (a, b) or field-collected and preserved at RT (c, d), and from *S. paniceum* specimens laboratory-reared and stored at -20°C (e, f) or field-collected and preserved at RT (g, h). **(B).** Representative thorax with legs MS spectra from *P. huttoni* (i, j), *O. ptilinoides* (k, l), *A. verbasci* (m, n), *X. rufovillosum* (o, p), and *A. punctatum* (q, r) field-collected and preserved at RT.

To assess the reproducibility and specificity, intra- and inter-species, respectively, of the MS spectra, MSP dendrogram were generated, taking into account the storing mode. In this way, three specimens per species and storing mode (frozenly or RT), were selected based on their intensity as representative samples. The first MSP dendrogram was created using uniquely Coleopteran MS spectra from laboratory reared adult specimens (frozenly stored), including the two species from the present study and the six species previously analysed (Additional Figure S1) (28). When the MS spectra of laboratory-reared *M. affine* and *S. paniceum* specimens were compared with those of Coleopteran species already included in the database, they formed well-defined clusters with short branch lengths. These results underlined a high reproducible and species-specificity of MS spectra between these eight Coleopteran species.

The addition of three representative MS spectra from remaining species of field collected specimens revealed that specimens of the same species clustering in the same branch, independently of the storing mode (Figure 3). The absence of intertwining between species indicated that several MS peaks should be shared among specimens of the same species whatever the storing mode, allowing to group them onto the MSP dendrogram. However, MS spectra from field-collected specimens stored at room temperature showed higher intra-species distance level of branch (ie, until more than two-fold), reflecting a greater spectral variability for this preservation mode, independently of the species.

**Figure 3.**
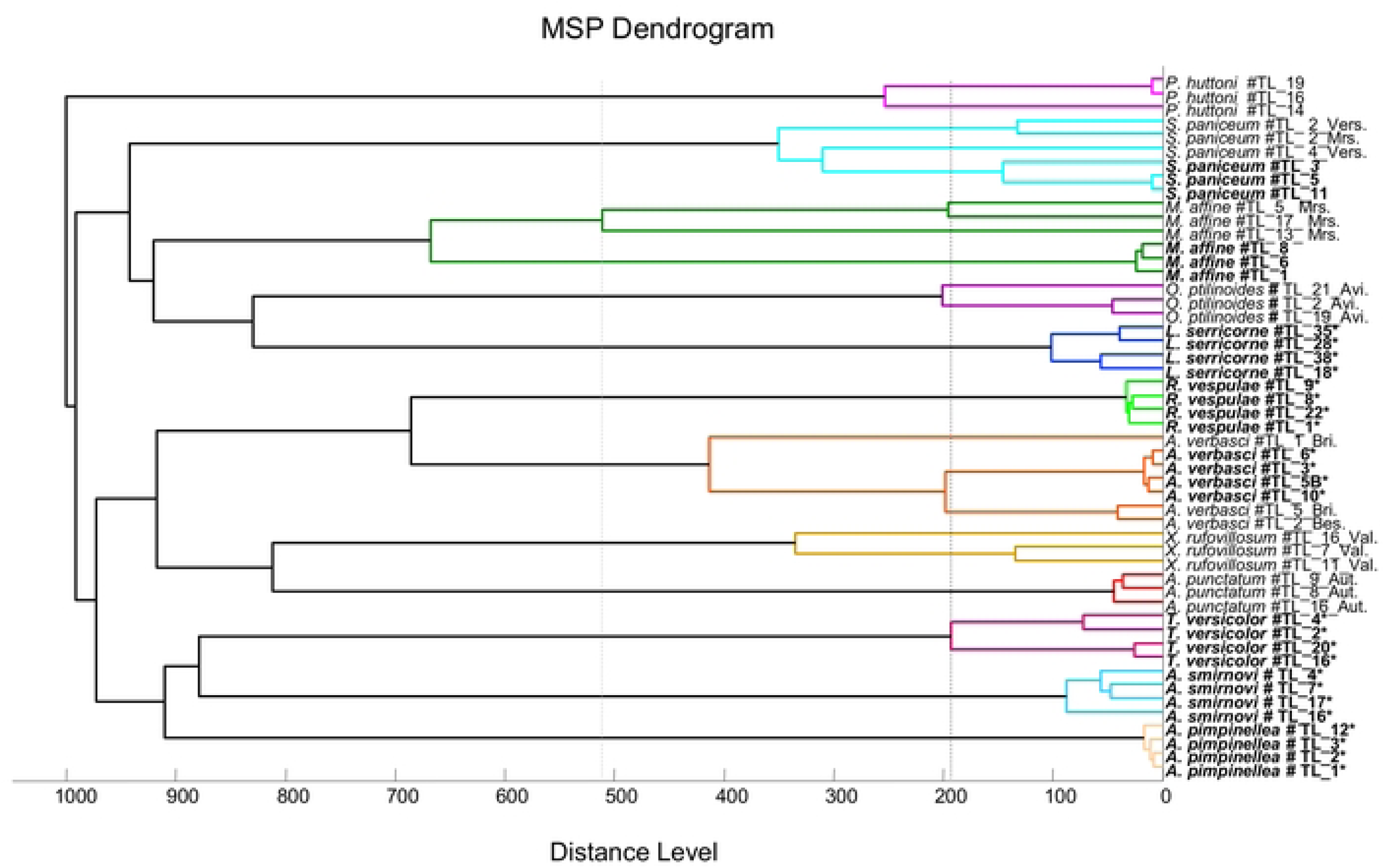
Assessment of species specificity of MS profiles using MSP Dendrogram from Coleopteran specimens originated from laboratory rearing and field collection. MSP dendrogram generated by Biotyper v3.0 software using three or four representative thorax and legs MS spectra of adult’s specimens. An arbitrary sample number, attributed to each specimen, is indicated after the species names (#TL_XX) followed by the site code for those field collected (Aut, Autain; Avi., Avignon; Bes., Besançon; Bri., Brive-la-Gaillarde; Mrs, Marseille; Vers., Versaille; Val., Valognes). Spectra included previously in our DB are indicated by asterisk (“*”) (DOI : <u>10.1016/j.ibiod.2024.106033</u>). Laboratory reared specimens are indicated in Bold. TL, thorax & legs. Black and grey dotted lines indicate the higher distance level of MS spectra among intra-species from specimens stored frozenly (laboratory-reared) and at room temperature (collected in the field), respectively.

**Figure 4.**
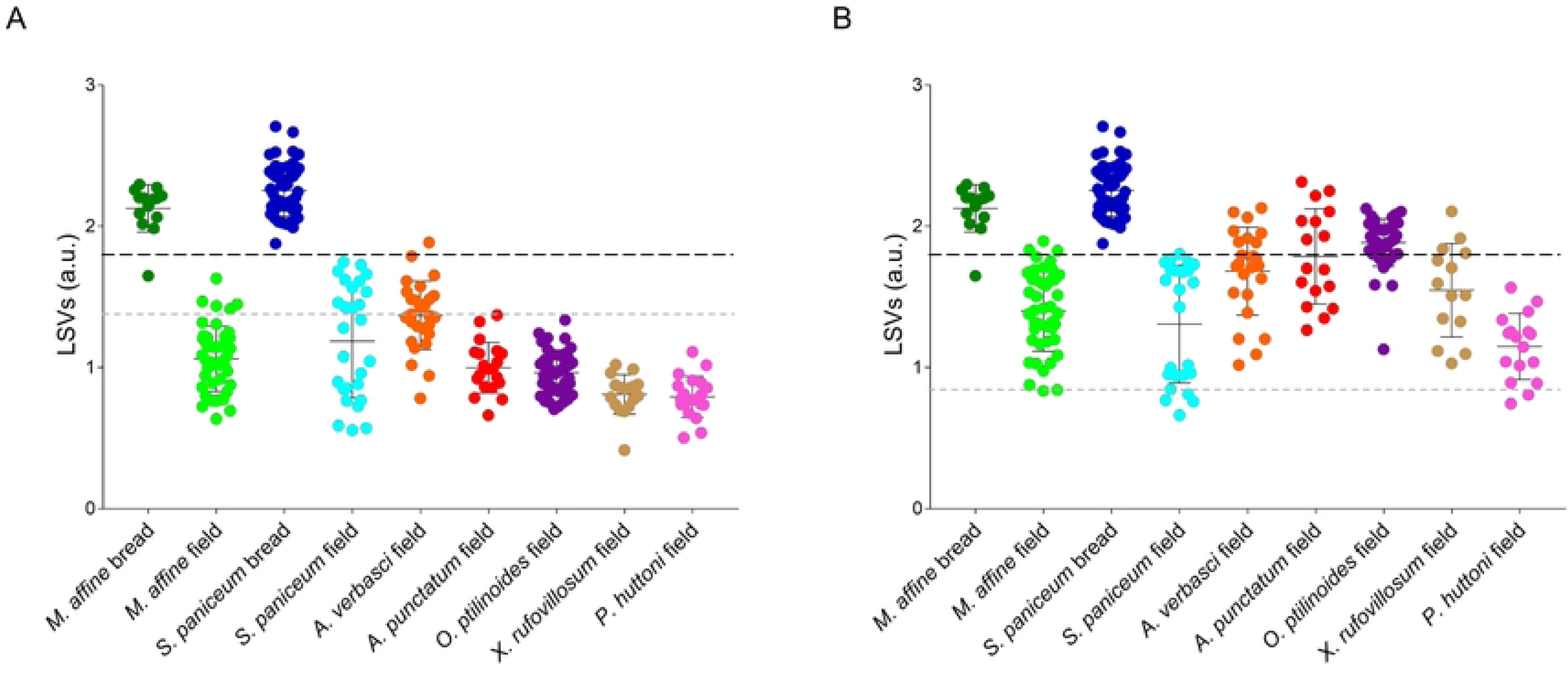
Evaluation of MALDI-TOF MS performances for Coleopteran species identification taking into account the specimen storing conditions and composition of the reference MS DB. **(A)** LSVs of thorax and legs MS spectra from adult Coleopteran specimens queried against the Database 1 (DB1). Among others, DB1 contains MS spectra from eight laboratory-reared Coleopteran species, for details see the paragraph “2.6” of materials and methods section. **(B)** LSVs of thorax and legs MS spectra from adult Coleopteran specimens queried against the Database 2 (DB2). DB2 is composed of the reference MS spectra included in the DB1 more those of the seven field-collected Coleopteran species from the present study (Table 1). The samples queried were indicated at the bottom of the graphic. The same color code was used per species and storing mode. The black dashed line represents the threshold value (LSV ≥ 1.8), for relevant identification. The grey dashed line represents the higher miss-identification LSV obtained for the dataset (panel A: 1.37, panel B: 0.85). a.u., arbitrary units; LSVs, log score values.

### 3.3. Blind tests

To assess the impact of specimen conservation conditions on the performances of the MALDI-TOF MS to identify correctly and relevantly Coleopteran at the species level, two successive blind tests were conducted. The first query was done against the DB1, a home-made DB of reference MS spectra from several arthropod species (44), including thorax and legs MS spectra from eight Coleopteran species coming uniquely from laboratory-rearing collection (Additional Figure S1). Among these eight Coleopteran species, two species (*M. affine* and *S. paniceum*) were newly added to the BD1, while *A. verbasci* was already included previously (28). Finally, no reference spectra were available in the DB1 for the other four species field collected (*A. punctatum, O. ptilinoides, P. huttoni and X. rufovillosum*,).

The query of the MS spectra from 267 specimens against the DB1, revealed that for both species originated from laboratory-rearing and frozenly stored (*M. affine* and *S. paniceum*), 100% of the samples were correctly identified. Moreover, the proportions of identification reliability (LSVs ≥ 1.8) were very high, with 92.9% (13/14) for *M. affine* and 100% (48/48) for *S. paniceum* (Table 2). For the three field collected species having homologous in the DB1, the rate of correct identification remained elevates, ranging from 89.1% (41/46) for *M. affine*, 92.6% (25/27) for *S. paniceum*, until 100% (27/27) for *A. verbasci*. Nevertheless, only one MS spectra from a *A. verbasci* specimen reach the threshold LSVs (≥ 1.8) to consider identification as relevant. For the four remaining species without homologous in the DB1, the higher miss-identification LSV not exceeding 1.4. The best value was obtained for a *A. punctatum* sample with a LSV of 1.37 (Figure 3A, Table 2). These low values indicated a weak spectral similarity between specimens stored frozenly and at RT.

To evaluate this last hypothesis, an upgrading of the DB1 by adding thorax and legs MS spectra from three specimens per species field collected was done to create the DB2 (Table 2). The 246 remaining MS spectra were query against the DB2 (Figure 3B). For laboratory-reared specimens from both species, identification results were unchanged by this upgrading, whereas for field collected specimens, the LSVs increased significantly for all species (Table 2, Wilcoxon matched-pairs signed rank test). Among the 184 MS spectra from specimens of field origin, concordant identification with morphological classification were obtained for 182 of them (98.9%). MS spectra from two *P. huttoni* were incorrectly classified and corresponded to the two lower LSVs for this group below 0.85 (Figure 3B). The proportion of correct and relevant (LSVs ≥ 1.8) identification varied according to species, ranging from 80.0% (36/45) for *O. ptilinoides* until 0.0% for *P. huttoni* (Table 2).

Interestingly, for *M. affine* and *S. paniceum*, the upgrading of the DB with MS spectra from field collected specimens did not succeeded to improve the proportion of correct and relevant identification of field samples, reaching 7.0% and 4.2%, respectively. Although for 33.3% and 47.1% of *A. verbasci* and *A. punctatum* MS spectra, respectively, reached the relevant threshold value (ie, ≥ 1.8), these proportions remain insufficient for its application to monitor these pest insects.

### 3.4. Assessment of geographical origin and storing time onto MS spectra

Among the seven Coleopteran species, uniquely for *A. verbasci* and *S. paniceum*, field collected specimens were available at different time points and geographical sites (Table 1). The classification of these two species according to the duration of storing and collection location was done and resulting LSVs were compared (Additional Figure S2). For *A. verbasci*, the LSVs appeared relatively homogeneous among collection sites and storing duration (Additional Figure S2A). This observation was objectified by the demonstration that no significant variation of LSVs (Kruskal-Wallis test, *p > 0.05*) was noticed between collection sites and dates of *A. verbasci* specimens. It is interesting to notice that for specimens collected in Versailles and stored ten years (ie, 2014) at RT, their LSVs are around of the threshold, suggesting a relative good storing of their protein profiles.

For *S. paniceum*, results were more dichotomous, with one half (n=16, 59.3%) presenting a LSV upper than 1.4 and another half (n=11, 40.7%) harbouring a LSV lower than 1 (Additional Figure S2B). As in several sites only one specimen was collected, statistical tests could not be done. However, the procurement of relatively high LSVs in several sites and distinct time points, notably for the older, suggested a relative stability of MS spectra independently of their geographical origin and/or the duration of samples storing at RT. Interestingly, the highest LSVs were obtained for the collection sites from which the reference MS spectra also originated for both species (Additional Figure S2A and S2B).

To obtain an overview of the intra- and inter-species MS spectra diversity, proteins profiles from field collection were compared to their laboratory reared counterpart species using CCI. These comparisons revealed that higher MS spectra correlation index were obtained for reared species, with 0.56 and 0.63 for *A. verbasci* and *S. paniceum*, respectively (Additional Figure S2B). These values fallen to 0.32 and 0.29 for these last two species field collected, underlining a reduction of intra-species homogeneity of MS spectra. The splicing of MS spectra from *S. paniceum* field collected according to their LSVs with a cut-off at 1.4, confirmed on the heatmap the weak correlation of these protein profiles with low LSVs (field^L^ group), among them (ie, 0.27), but also compared to field specimens with LSVs above 1.4 (ie, field^H^ group). These results underlined that the low LSVs obtained for field^L^ group was attributed to the high intra-species diversity far from reference MS spectra. Paired species comparisons of MS spectra according to storing mode revealed that the levels of MS spectra correlation (0.32 and 0.28 for *A. verbasci* and *S. paniceum*) were similar as those obtained for field intra-species analyses.

### 3.5. Assessment of molecular tools for identification Coleopteran specimens according to their storing mode

As molecular identification is relatively time-consuming and expensive, only the specimens included in the MS database as references were submitted to COI and 16S target gene amplification and sequencing. The abdomens of three adult individuals from each of the seven selected species were processed, resulting in a total of 21 samples (Table 3). Moreover, three reared species, *M. affine* and *S. paniceum*, were used as positive controls to validate the molecular protocol. Amplification and sequencing success rates were assessed for both these reared species and the field-collected specimens. For *M. affine* of reared origin, amplification and sequencing were successful for both 16S and COI markers. The 16S sequences showed 98– 100% coverage with 92–95% identity, while COI sequences reached 97% coverage and 89% identity. For *S. paniceum* of reared origin, only the COI marker was successfully amplified and sequenced, yielding 100% coverage and identity, whereas the 16S marker failed to amplify. For field species, amplification with the 16S marker was successful for only four species (*A. verbasci*, *M. affine*, *O. ptilinoides*, and *P. huttoni*), with a PCR success rate of 42.9% (n=9/21) and a sequencing success rate of reaching solely 9.5% (n=2/21). For the COI marker, amplification of field samples was successful for four species (*A. verbasci, P. huttoni, S. paniceum* and *X. rufovillosum*), with a PCR success rate of 33.3% and a sequencing success rate of 9.5% (n=2/21). Despite successful amplification and sequencing in some cases, the resulting coverage was occasionally low (reaching as little as 27%) due to poor sequence quality.

**Table 3.** Comparison of molecular analysis effectiveness on abdomen samples per Coleopteran species and origins.

|  |  | 16S |  |  |  |  | COI |  |  |  |  |
| --- | --- | --- | --- | --- | --- | --- | --- | --- | --- | --- | --- |
| Species | Origin | Number of PCR success (%) | Number of sequencing success (%) | Accession number (NCBI) | Percent of coverage (Range) | Percent of identity (Range) | Number of PCR success (%) | Number of sequencing success (%) | Accession number (NCBI) | Percent of coverage (Range) | Percent of identity (Range) |
| *Ano. punctatum* | Field (n=3) | 0 (-) | 0 (-) | (-) | (-) | (-) | 0 (-) | 0 (-) | (-) | (-) | (-) |
| *Ant. verbasci* | Field (n=3) | 2 (66.6%) | 2 (66.6%) | KX087239.1 | [27- 60] | [83-95] | 2 (66.6%) | 0 (-) | (-) | (-) | (-) |
| *M. affine* | Bread (n=3) | 3 (100%) | 2 (66.6%) | JN097759.1 | [98-100] | [92-95] | 3 (100%) | 1 (33.3%) | KU494140.1 | [97] | [89] |
|  | Field (n=3) | 3 (100%) | 0 (-) | (-) | (-) | (-) | 0 (-) | 0 (-) | (-) | (-) | (-) |
| *O. ptilinoides* | Field (n=3) | 2 (66.6%) | 0 (-) | (-) | (-) | (-) | 0 (-) | 0 (-) | (-) | (-) | (-) |
| *P. huttoni* | Field (n=3) | 2 (66.6%) | 0 (-) | (-) | (-) | (-) | 1 (33.3%) | 0 (-) | (-) | (-) | (-) |
| *S. paniceum* | Bread (n=3) | 0 (-) | 0 (-) | (-) | (-) | (-) | 3 (100%) | 1 (33.3%) | MH817140.1 | [100] | [100] |
|  | Field (n=3) | 0 (-) | 0 (-) | (-) | (-) | (-) | 3 (100 %) | 2 (66.6%) | MH817140.1 | [100] | [99.5] |
| *X. rufovillosum* | Field (n=3) | 0 (-) | 0 (-) | (-) | (-) | (-) | 1 (33.3%) | 0 (-) | (-) | (-) | (-) |
| **Total n (%)** * | **27** | **12 (44.4%)** | **4 (14.8%)** |  |  |  | **13 (48.1%)** | **4 (14.8%)** |  |  |  |

\*The proportion among the total number tested, n=27. n, number, *Ano.*, *Anobium*; *Ant.*, *Anthrenus*; *M.*, *Mezium*; *O.*, *Oligomerus*; *P.*, *Pentarthrum*; *S.*, *Stegobium*; *X.*, *Xestobium*.

## 4. Discussion

Over the past decade, MALDI-TOF MS has emerged as a powerful tool for the identification of arthropods of medical and veterinary importance (45). Numerous studies have demonstrated the robustness of MALDI-TOF MS for the accurate identification of a wide range of arthropods, distinguishing vector from non-vector species (46,47). This innovative approach contributed to improve vector monitoring in real-time at low cost (46,48) and has also proven effective in detecting the infectious status of certain arthropod vectors (49). More recently, MALDI-TOF MS has been successfully applied to the classification of insect pests damaging cultural heritage (28) offering new perspectives for the monitoring and protection of museum collections (50). This pioneering study focused on eight laboratory-reared Coleoptera species at various developmental stages (i.e., adult, larva, and exuviae). Here, seven Coleoptera species at adult stage collected essentially in museums were included, focusing on the impact of conservation conditions and geographical origin on spectral profiles.

For the two laboratory-reared species tested (*ie*, *M. affine, S. paniceum*) and frozenly stored, the high reproducibility and species-specificity of MS spectra confirmed MALDI-TOF MS efficiency for correct and reliable identification with rate reaching 98.4% (61/62). For the other specimens field collected with homolog in the MS DB, although correct identification was obtained, only one *A. verbasci* specimen was considered relevantly identified. The reliability of specimen identification by MS depends on their inter-species specificity, but also on their reproducibility within a given species (51). Several factors can affect the reproducibility and intensity of MS spectra from specimens of the same arthropod species. These factors can be separated into two categories, those depending on intrinsic factors related to the sample and those on extrinsic factors linked to environmental conditions (52). Regarding intrinsic factors, these mainly involve changes in protein expression profiles related to the tissue compartment and/or developmental stage selected (30,53,54).The extrinsic factors include, among others, the duration and the storing mode (31,55), the conditions of sample preparation (31,56) and the geographical origin of the samples (57), which are also other parameters impairing the intra-species repeatability of MS profiles. Here, field collected specimens preserved at RT for 1 to 10 years were reliably identified by MALDI-TOF MS only when homologous reference spectra stored in the same conditions were available in the database; in their absence, identification scores remained low. These low values reflect weak spectral similarity between specimens preserved under different conditions (frozen vs. room temperature), underscoring the limitations of this proteomic approach when no matching reference is present. Our findings indicate that sample identification by MALDI-TOF MS was significantly increased when the database contains spectra from homologous species preserved under comparable conditions. These results align with several studies highlighting the crucial role of preservation methods and storage duration on spectral quality, but also the content of the reference MS DB (58). For instance, high MS identification accuracy (up to 99.6%) was obtained for ticks field collected and preserved either in ethanol or frozenly stored on the condition that the reference database included spectra from specimens preserved in similar circumstances (59). Nevertheless, it is not always sufficient. More recently, Saidou *et al.* demonstrated that long-term (ie, upper than ten years) ethanol preservation of ticks negatively impacted MALDI-TOF MS identification accuracy, impairing matching with homolog species stored in the same conditions during few years (60). An optimization of the protocol was then required for the creation of new reference spectra to enhance their identification. It is probable that for museum collections of Coleopteran stored in RT during decades, a degradation of protein profile occurred, underlying the same strategy(61).

Here, for the two Coleopteran species preserved at RT over a short period (about one year) to a long period (up to ten years), intra-species MS profiles were considered as relatively stable independently of storing time, after exclusion of outlier spectra of low quality from *P. paniceum*. Moreover, the absence of significant variation of identification score throughout the time for *A. verbasci*, further supported the reproducibility. This observation was supported by previous studies, demonstrating that although sample MS profiles undergo rapid modification shortly after preservation depending on the storage method (eg, ethanol, RT…), compared to fresh or frozen counter species samples, a stabilization followed by reproducibility of MS spectra was noticed thereafter few months of storing (62). It would be interesting to determine the delay necessary of storing at RT for some Coleopteran species to obtain stable and reproducible MS spectra. These findings highlight the robustness of MALDI-TOF MS for arthropod identification, even with extended preservation times, provided that initial handling was carefully controlled.

In the present work, the low rate of identification considered reliable for field specimens could be attributed to high background noise in the spectra. Indeed, this high background noise generated intra-species heterogeneity hindered the generation of high LSVs. Since the LSV calculation depends on both shared and unique peaks between the query and reference spectra, spectral reproducibility and diversity are critical parameters. To limit inaccurate identification, MS spectra with a maximum peak intensity below 3,000 a.u. are generally considered as non-compliant and are then excluded from the analysis (63). This maximum peak intensity cut-off was established either for arthropod samples preserved frozen or in alcohol, limiting protein degradation compared to a preservation at RT (64). Long-term storage at RT of the Coleopterans have likely participated to the integrity loss of species-specific proteins inducing spectra intensity decrease, profile heterogeneity and high background noise. As long-term preservation of arthropod sample, generally induced decrease of protein profile intensity, such MS spectra conformity criteria was not applied to prevent in priority the exclusion spectra stored during long time independently of their reproducibility.

Moreover, the low quality of some spectra was attributed to the duration of storing, but the unknown interval between the death of the specimens and their collection in the field. These uncertain post-mortem intervals should also participate to protein degradation and suboptimal preservation conditions. Indeed, several studies have reported that delays between specimen collection and processing, as well as prolonged storage at ambient or elevated temperatures, significantly compromise protein integrity, resulting in altered and lower-quality of protein spectra (65–67).

It is interesting to note that *P. huttoni* samples generated spectra of low diversity. This low diversity could explain in part the failing of correct and relevant identification obtained for this group. It would be valuable to determine, in the future, the minimum number and intensity of MS peaks required in reference and query spectra to ensure reliable spectral matching.

Finally, in the absence of entomological expertize or in cases of taxonomic uncertainty, target gene sequencing using molecular biology techniques was frequently employed for accurate identification of arthropod species (68,69). Here, uniquely Coleopteran specimens added to the reference MS spectra DB were submitted to molecular biology to confirm morphological identification. Overall, a very low sequencing success rate (<15%) was obtained for both target genes (ie, 16S and COI). The long-term storage (i.e., more than one year) of the specimens at ambient temperature have probably compromised DNA integrity. The detrimental effects of specimen ambient storage on DNA quality have been repeatedly demonstrated on arthropods (PMID: 31990210). For instance, ant specimens preserved at RT during 10 months exhibited a substantial loss of amplifiable DNA, with PCR success rates dropping below 50% (70). Moreover, the unknown interval time between Coleopteran death and it field collection is another period of suboptimal storage during which DNA fragmentation occurs (71). Nevertheless, the poor sequencing performance observed even in fresh specimens is questioning on the application of the appropriate DNA extraction method or the reagents used for amplification and PCR product sequencing (72,73). Previous studies reported that even with fresh Coleopteran specimens, molecular identification could be challenging (28,74,75).

Secondly, the frequent occurrence of taxonomic inaccuracies within public DNA sequence databases (76) undermines the reliability of species-level identification. These errors are often compounded by missing or incomplete metadata, such as specimen origin, preservation method, or collection date (77). Such gaps in data can hinder accurate comparisons and reduce the interpretability of sequence matches, particularly when dealing with degraded or poorly preserved specimens, as in the present study. Taken together, these findings reinforce the conclusion that inadequate storage conditions can severely compromise both the effectiveness of molecular identification techniques and the quality and reproducibility of MS spectra.

## 5. Conclusion

The rapid and accurate identification of pest insects is essential for effective control strategies. In this context, MALDI-TOF MS proved to be a powerful and reliable alternative to traditional and molecular methods, particularly for fresh specimens. Its performance remains superior even when field-collected samples are sub optimally preserved, whereas molecular techniques often fail due to DNA degradation and low amplification success. Morphological identification, though informative, requires extensive taxonomic expertise and is not easily scalable. The success of MALDI-TOF MS increases when reference spectra from similarly preserved specimens are available in the database. To optimize identification accuracy, improving post-collection preservation—using ethanol or desiccants—is crucial. Additionally, reducing the time between sampling and analysis limits protein and DNA degradation. Future efforts should prioritize immediate analysis of field specimens to ensure high-quality identification results.

## List of abbreviations

CHCA: α-cyano-4-hydroxycinnamic acid.
CICRP: Centre Interdisciplinaire de Conservation et de Restauration du Patrimoine.
COI: cytochrome oxidase subunit I.
DNA: Deoxyribonucleic Acid.
HPLC: High-Performance Liquid Chromatography.
ITS: Internal transcribed spacer
LSV: log score value.
MALDI-TOF MS: Matrix-Assisted Laser Desorption/Ionization Time-of-Flight Mass Spectrometry.
PCA: Principal Component Analysis.
PCR: Polymerase Chain Reaction.
RT: Room temperature.

## Availability of data and materials

The MS reference spectra included in the database of this study are freely accessible and can be downloaded via the provided DOI.

## Conflict of interest

The authors declare that there are no conflicts of interest.

## Ethics approval and consent to participate

Not applicable.

## Consent for publication

Not applicable.

## Funding

This work has been supported by the *Délégation Générale pour l’Armement* (DGA, MSProfileR project, Grant no PDH-2-NBC 2-B-2201).

This work was supported by the Institut IHU Méditerrannée Infection. DH received a PhD grant from the IHU Méditerrannée Infection. Funding played no role in the study design, data collection and analysis, decision to publish or preparation of the manuscript.

## Authors’ contributions

Conceived and designed the experiments: LA

Performed the experiments: DH, LA

Analyzed the data: DH, LA

Contributed reagents/materials/analysis tools: DH, AZD, JMB, PP, FF

Investigation: DH, FF, JMB

Drafted the paper: DH, LA, PP

Revised critically the paper: all the authors.

## Additional files

**Additional Figure S1. Assessment of species specificity of MS profiles using MSP dendrogram from laboratory-reared Coleopteran specimens.** MSP dendrogram generated by Biotyper v3.0 software using three or four representative thorax and legs MS spectra of adult’s specimens originated from laboratory rearing. An arbitrary sample number, attributed to each specimen, is indicated after the species names (#TL_XX). Spectra included previously in our DB are indicated by asterisk (“*”) (DOI : <u>10.1016/j.ibiod.2024.106033</u>). TL, thorax & legs.

**Additional Figure S2. Consequences of storing mode and duration onto Coleopteran MS spectra. (A)** LSVs of thorax and legs MS spectra from adult *A. verbasci* specimens collected in the field and queried against the Database 2 (DB2). **(B)** LSVs of thorax and legs MS spectra from adult *P. paniceum* specimens collected in the field and queried against the Database 2 (DB2). The collection site and year of sampling were indicated at the bottom of the graphic. All these specimens were stored at RT. The black dashed line represents the threshold value (LSV ≥ 1.8), for relevant identification. *Reference spectra coming from the same collection site. a.u., arbitrary units; LSVs, log score values. **(C)** Composite Correlation Index (CCI) matrix value representing the levels of MS spectra reproducibility among specimens of the same species and between species (*A. verbasci*, *P. paniceum*), according to storing mode and duration. MS spectra from field collected specimens plus twenty specimens per species laboratory reared were also included in this analysis. The levels of MS spectra reproducibility are indicated in red and blue, revealing relatedness and incongruence between spectra, respectively. The values correspond to the mean coefficient correlation and respective standard deviations obtained for paired condition comparisons. Field^H/L^, classification of samples according to their high (H) or low (L) LSVs. CCI was calculated with MALDI-Biotyper v.3.0 software.

## Notes

### Competing Interest Statement

The authors have declared no competing interest.

## References

1. Gorman GE, Shep SJ. Preservation Management for Libraries, Archives and Museums [Internet]. Facet Publishing. 2006 [cited 2025 Jul 29]. Available from: https://www.routledge.com/Preservation-Management-for-Libraries-Archives-and-Museums/Gorman-Shep/p/book/9781856045742

2. Magali Toriti, Aline Durand & Fabien Fohrer. Traces d’insectes xylophages communs dans le bois : Atlas d’identification - Europe occidentale | SpringerLink [Internet]. Springer International Publishing. [cited 2025 Jul 29]. Available from: https://link.springer.com/book/10.1007/978-3-030-66391-9

3. Konsa K, Kormpaki T, Turu J. Biological Damage to Textiles and Prevention Methods. In: Jose S, Thomas S, Pandit P, Pandey R, editors. Handbook of Museum Textiles [Internet]. 1st ed. Wiley; 2022 [cited 2025 Jul 29]. p. 23–43. Available from: https://onlinelibrary.wiley.com/doi/10.1002/9781119983439.ch2

4. Brimblecombe P, Rohrer U, Landsberger B, Querner P. Insect catch at historic libraries in rural and urban settings. Int Biodeterior Biodegrad [Internet]. 2024 Aug [cited 2025 Jul 29];193:105855. Available from: https://linkinghub.elsevier.com/retrieve/pii/S0964830524001264

5. Ignatowicz S, Janczukowicz K, Olejarski P. Integrated Pest Management (IPM) of the drug store beetle, Stegobium paniceum (L.), a serious pest of old books. J Entomol Acarol Res [Internet]. 2011 Aug 20 [cited 2025 Jul 29];43(2):177. Available from: http://www.pagepressjournals.org/index.php/jear/article/view/jear.2011.177

6. Cao Y, Pistillo OM, Lou Y, D’Isita I, Maggi F, Hu Q, et al. Electrophysiological and behavioural responses of Stegobium paniceum to volatile compounds from Chinese medicinal plant materials. Pest Manag Sci. 2022 Aug;78(8):3697–703.

7. Querner P. Insect Pests and Integrated Pest Management in Museums, Libraries and Historic Buildings. Insects. 2015 Jun 16;6(2):595–607.

8. Dara SK. The New Integrated Pest Management Paradigm for the Modern Age. J Integr Pest Manag [Internet]. 2019 Jan 1 [cited 2025 Jul 29];10(1). Available from: https://academic.oup.com/jipm/article/doi/10.1093/jipm/pmz010/5480541

9. Ferret-Bouin P. Clé illustrée des familles des coléoptères de France. L’Entomologiste. 1995;7((3)):45–52.

10. Barrett RDH, Hebert PDN. Identifying spiders through DNA barcodes. Can J Zool [Internet]. 2005 Mar 1 [cited 2025 Jul 29];83(3):481–91. Available from: http://www.nrcresearchpress.com/doi/10.1139/z05-024

11. Murugan K, Vadivalagan C, Karthika P, Panneerselvam C, Paulpandi M, Subramaniam J, et al. DNA barcoding and molecular evolution of mosquito vectors of medical and veterinary importance. Parasitol Res. 2016 Jan;115(1):107–21.

12. Trematerra P, Pinniger D. Museum Pests–Cultural Heritage Pests. In: Athanassiou CG, Arthur FH, editors. Recent Advances in Stored Product Protection [Internet]. Berlin, Heidelberg: Springer Berlin Heidelberg; 2018 [cited 2025 Jul 29]. p. 229–60. Available from: http://link.springer.com/10.1007/978-3-662-56125-6_11

13. Tautz D, Arctander P, Minelli A, Thomas RH, Vogler AP. A plea for DNA taxonomy. Trends Ecol Evol [Internet]. 2003 Feb [cited 2025 Jul 29];18(2):70–4. Available from: https://linkinghub.elsevier.com/retrieve/pii/S0169534702000411

14. Nagoshi RN, Brambila J, Meagher RL. Use of DNA Barcodes to Identify Invasive Armyworm Spodoptera Species in Florida. J Insect Sci [Internet]. 2011 Nov [cited 2025 Jul 29];11(154):1–11. Available from: https://academic.oup.com/jinsectscience/article-lookup/doi/10.1673/031.011.15401

15. Taberlet P, Coissac E, Pompanon F, Brochmann C, Willerslev E. Towards next-generation biodiversity assessment using DNA metabarcoding. Mol Ecol [Internet]. 2012 Apr [cited 2025 Jul 29];21(8):2045–50. Available from: https://onlinelibrary.wiley.com/doi/10.1111/j.1365-294X.2012.05470.x

16. Hebert PDN, Ratnasingham S, De Waard JR. Barcoding animal life: cytochrome c oxidase subunit 1 divergences among closely related species. Proc R Soc Lond B Biol Sci [Internet]. 2003 Aug 7 [cited 2025 Jul 29];270(suppl_1). Available from: https://royalsocietypublishing.org/doi/10.1098/rsbl.2003.0025

17. Steinke D, Vences M, Salzburger W, Meyer A. TaxI: a software tool for DNA barcoding using distance methods. Philos Trans R Soc Lond B Biol Sci. 2005 Oct 29;360(1462):1975–80.

18. Bridge PD, Roberts PJ, Spooner BM, Panchal G. On the unreliability of published DNA sequences. New Phytol. 2003 Oct;160(1):43–8.

19. Nilsson RH, Ryberg M, Kristiansson E, Abarenkov K, Larsson KH, Kõljalg U. Taxonomic Reliability of DNA Sequences in Public Sequence Databases: A Fungal Perspective. Fairhead C, editor. PLoS ONE [Internet]. 2006 Dec 20 [cited 2025 Jul 29];1(1):e59. Available from: https://dx.plos.org/10.1371/journal.pone.0000059

20. Floyd R, Lima J, deWaard J, Humble L, Hanner R. Common goals: policy implications of DNA barcoding as a protocol for identification of arthropod pests. Biol Invasions [Internet]. 2010 Sep [cited 2025 Jul 29];12(9):2947–54. Available from: http://link.springer.com/10.1007/s10530-010-9709-8

21. Porter TM, Hajibabaei M. Scaling up: A guide to high-throughput genomic approaches for biodiversity analysis. Mol Ecol. 2018 Jan;27(2):313–38.

22. Tedersoo L, Drenkhan R, Anslan S, Morales-Rodriguez C, Cleary M. High-throughput identification and diagnostics of pathogens and pests: Overview and practical recommendations. Mol Ecol Resour. 2019 Jan;19(1):47–76.

23. Querner P, Szucsich N, Landsberger B, Brimblecombe P. DNA Metabarcoding Analysis of Arthropod Diversity in Dust from the Natural History Museum, Vienna. Diversity [Internet]. 2024 Aug 6 [cited 2025 Jul 29];16(8):476. Available from: https://www.mdpi.com/1424-2818/16/8/476

24. Beng KC, Tomlinson KW, Shen XH, Surget-Groba Y, Hughes AC, Corlett RT, et al. The utility of DNA metabarcoding for studying the response of arthropod diversity and composition to land-use change in the tropics. Sci Rep. 2016 Apr 26;6:24965.

25. Elbrecht V, Vamos EE, Meissner K, Aroviita J, Leese F. Assessing strengths and weaknesses of DNA metabarcoding-based macroinvertebrate identification for routine stream monitoring. Yu D, editor. Methods Ecol Evol [Internet]. 2017 Oct [cited 2025 Jul 29];8(10):1265–75. Available from: https://besjournals.onlinelibrary.wiley.com/doi/10.1111/2041-210X.12789

26. Piper AM, Batovska J, Cogan NOI, Weiss J, Cunningham JP, Rodoni BC, et al. Prospects and challenges of implementing DNA metabarcoding for high-throughput insect surveillance. GigaScience. 2019 Aug 1;8(8):giz092.

27. Batovska J, Piper AM, Valenzuela I, Cunningham JP, Blacket MJ. Developing a non-destructive metabarcoding protocol for detection of pest insects in bulk trap catches. Sci Rep. 2021 Apr 12;11(1):7946.

28. Hamadouche D, Diarra AZ, Fohrer F, Bérenger JM, Benakhla A, Almeras L, et al. Assessment of MALDI-TOF MS for the identification of cultural heritage insect pests. Int Biodeterior Biodegrad [Internet]. 2025 Mar [cited 2025 Jul 29];199:106033. Available from: https://linkinghub.elsevier.com/retrieve/pii/S096483052500037X

29. Karger A, Kampen H, Bettin B, Dautel H, Ziller M, Hoffmann B, et al. Species determination and characterization of developmental stages of ticks by whole-animal matrix-assisted laser desorption/ionization mass spectrometry. Ticks Tick-Borne Dis. 2012 Apr;3(2):78–89.

30. Nabet C, Kone AK, Dia AK, Sylla M, Gautier M, Yattara M, et al. New assessment of Anopheles vector species identification using MALDI-TOF MS. Malar J. 2021 Jan 9;20(1):33.

31. Nebbak A, Willcox AC, Bitam I, Raoult D, Parola P, Almeras L. Standardization of sample homogenization for mosquito identification using an innovative proteomic tool based on protein profiling. Proteomics. 2016 Dec;16(24):3148–60.

32. Yssouf A, Almeras L, Raoult D, Parola P. Emerging tools for identification of arthropod vectors. Future Microbiol. 2016;11(4):549–66.

33. Costa MM, Guidez A, Briolant S, Talaga S, Issaly J, Naroua H, et al. Identification of Neotropical Culex Mosquitoes by MALDI-TOF MS Profiling. Trop Med Infect Dis. 2023 Mar 13;8(3):168.

34. Kumsa B, Laroche M, Almeras L, Mediannikov O, Raoult D, Parola P. Morphological, molecular and MALDI-TOF mass spectrometry identification of ixodid tick species collected in Oromia, Ethiopia. Parasitol Res. 2016 Nov;115(11):4199–210.

35. Benkacimi L, Gazelle G, El Hamzaoui B, Bérenger JM, Parola P, Laroche M. MALDI-TOF MS identification of Cimex lectularius and Cimex hemipterus bedbugs. Infect Genet Evol J Mol Epidemiol Evol Genet Infect Dis. 2020 Nov;85:104536.

36. Ahamada M’madi S, Diarra AZ, Almeras L, Parola P. Identification of ticks from an old collection by MALDI-TOF MS. J Proteomics. 2022 Jul 30;264:104623.

37. Hamlili FZ, Thiam F, Laroche M, Diarra AZ, Doucouré S, Gaye PM, et al. MALDI-TOF mass spectrometry for the identification of freshwater snails from Senegal, including intermediate hosts of schistosomes. PLoS Negl Trop Dis. 2021 Sep;15(9):e0009725.

38. Bamou R, Costa MM, Diarra AZ, Martins AJ, Parola P, Almeras L. Enhanced procedures for mosquito identification by MALDI-TOF MS. Parasit Vectors. 2022 Jun 30;15(1):240.

39. Kumsa B, Laroche M, Almeras L, Mediannikov O, Raoult D, Parola P. Morphological, molecular and MALDI-TOF mass spectrometry identification of ixodid tick species collected in Oromia, Ethiopia. Parasitol Res. 2016 Nov;115(11):4199–210.

40. Folmer O, Black M, Hoeh W, Lutz R, Vrijenhoek R. DNA primers for amplification of mitochondrial cytochrome c oxidase subunit I from diverse metazoan invertebrates. Mol Mar Biol Biotechnol. 1994 Oct;3(5):294–9.

41. Olson RLO, Farris RE, Barr NB, Cognato AI. Molecular identification of Trogoderma granarium (Coleoptera: Dermestidae) using the 16s gene. J Pest Sci [Internet]. 2014 Dec [cited 2025 Jul 29];87(4):701–10. Available from: http://link.springer.com/10.1007/s10340-014-0621-3

42. Olson RLO, Farris RE, Barr NB, Cognato AI. Molecular identification of Trogoderma granarium (Coleoptera: Dermestidae) using the 16s gene. J Pest Sci [Internet]. 2014 Dec [cited 2025 Jul 29];87(4):701–10. Available from: http://link.springer.com/10.1007/s10340-014-0621-3

43. Benson DA, Cavanaugh M, Clark K, Karsch-Mizrachi I, Lipman DJ, Ostell J, et al. GenBank. Nucleic Acids Res. 2013 Jan;41(Database issue):D36–42.

44. Yssouf A, Almeras L, Raoult D, Parola P. Emerging tools for identification of arthropod vectors. Future Microbiol. 2016;11(4):549–66.

45. Sevestre J, Diarra AZ, Laroche M, Almeras L, Parola P. Matrix-assisted laser desorption/ionization time-of-flight mass spectrometry: an emerging tool for studying the vectors of human infectious diseases. Future Microbiol. 2021 Mar;16:323–40.

46. Flacio E, Engeler L, Tonolla M, Lüthy P, Patocchi N. Strategies of a thirteen year surveillance programme on Aedes albopictus (Stegomyia albopicta) in southern Switzerland. Parasit Vectors. 2015 Apr 9;8:208.

47. Vega-Rúa A, Pagès N, Fontaine A, Nuccio C, Hery L, Goindin D, et al. Improvement of mosquito identification by MALDI-TOF MS biotyping using protein signatures from two body parts. Parasit Vectors. 2018 Nov 3;11(1):574.

48. Nebbak A, Willcox AC, Koumare S, Berenger JM, Raoult D, Parola P, et al. Longitudinal monitoring of environmental factors at Culicidae larval habitats in urban areas and their association with various mosquito species using an innovative strategy. Pest Manag Sci. 2019 Apr;75(4):923–34.

49. Sevestre J, Diarra AZ, Laroche M, Almeras L, Parola P. Matrix-assisted laser desorption/ionization time-of-flight mass spectrometry: an emerging tool for studying the vectors of human infectious diseases. Future Microbiol. 2021 Mar;16:323–40.

50. Venette RC, Hutchison WD. Invasive Insect Species: Global Challenges, Strategies & Opportunities. Front Insect Sci. 2021;1:650520.

51. Yssouf A, Almeras L, Raoult D, Parola P. Emerging tools for identification of arthropod vectors. Future Microbiol. 2016;11(4):549–66.

52. Bouledroua R, Hamadouche D, Amirat Z, Alméras L. Empreintes de spectrométrie de masse appliquées à l’étude des culicidés. Rev Francoph Lab [Internet]. 2025 Mar [cited 2025 Jul 29];2025(570):34–46. Available from: https://linkinghub.elsevier.com/retrieve/pii/S1773035X25000188

53. Schaffner F, Kaufmann C, Pflüger V, Mathis A. Rapid protein profiling facilitates surveillance of invasive mosquito species. Parasit Vectors. 2014 Mar 31;7:142.

54. Nebbak A, Koumare S, Willcox AC, Berenger JM, Raoult D, Almeras L, et al. Field application of MALDI-TOF MS on mosquito larvae identification. Parasitology. 2018 Apr;145(5):677–87.

55. Nebbak A, El Hamzaoui B, Berenger JM, Bitam I, Raoult D, Almeras L, et al. Comparative analysis of storage conditions and homogenization methods for tick and flea species for identification by MALDI-TOF MS. Med Vet Entomol. 2017 Dec;31(4):438–48.

56. Diarra AZ, Almeras L, Laroche M, Berenger JM, Koné AK, Bocoum Z, et al. Molecular and MALDI-TOF identification of ticks and tick-associated bacteria in Mali. PLoS Negl Trop Dis. 2017 Jul;11(7):e0005762.

57. Bamou R, Costa MM, Diarra AZ, Martins AJ, Parola P, Almeras L. Enhanced procedures for mosquito identification by MALDI-TOF MS. Parasit Vectors. 2022 Jun 30;15(1):240.

58. Costa MM, Corbel V, Ben Hamouda R, Almeras L. MALDI-TOF MS Profiling and Its Contribution to Mosquito-Borne Diseases: A Systematic Review. Insects. 2024 Aug 29;15(9):651.

59. Diarra AZ, Almeras L, Laroche M, Berenger JM, Koné AK, Bocoum Z, et al. Molecular and MALDI-TOF identification of ticks and tick-associated bacteria in Mali. PLoS Negl Trop Dis. 2017 Jul;11(7):e0005762.

60. Ahamada M’madi S, Diarra AZ, Almeras L, Parola P. Identification of ticks from an old collection by MALDI-TOF MS. J Proteomics. 2022 Jul 30;264:104623.

61. Ahamada M’madi S, Diarra AZ, Almeras L, Parola P. Identification of ticks from an old collection by MALDI-TOF MS. J Proteomics. 2022 Jul 30;264:104623.

62. Benyahia H, Parola P, Almeras L. Evolution of MALDI-TOF MS Profiles from Lice and Fleas Preserved in Alcohol over Time. Insects. 2023 Oct 20;14(10):825.

63. Hamlili FZ, Thiam F, Laroche M, Diarra AZ, Doucouré S, Gaye PM, et al. MALDI-TOF mass spectrometry for the identification of freshwater snails from Senegal, including intermediate hosts of schistosomes. Almeida IC, editor. PLoS Negl Trop Dis [Internet]. 2021 Sep 13 [cited 2025 Jul 29];15(9):e0009725. Available from: https://dx.plos.org/10.1371/journal.pntd.0009725

64. Topić Popović N, Kazazić SP, Bojanić K, Strunjak-Perović I, Čož-Rakovac R. Sample preparation and culture condition effects on MALDI-TOF MS identification of bacteria: A review. Mass Spectrom Rev [Internet]. 2023 Sep [cited 2025 Jul 29];42(5):1589–603. Available from: https://analyticalsciencejournals.onlinelibrary.wiley.com/doi/10.1002/mas.21739

65. Reeve MA, Buddie AG. A simple and inexpensive method for practical storage of field-sample proteins for subsequent MALDI-TOF MS analysis. Plant Methods. 2018;14:90.

66. Zissler A, Stoiber W, Steinbacher P, Geissenberger J, Monticelli FC, Pittner S. Postmortem Protein Degradation as a Tool to Estimate the PMI: A Systematic Review. Diagn Basel Switz. 2020 Nov 26;10(12):1014.

67. Chhikara A, Kumari P, Dalal J, Kumari K. Protein degradation patterns as biomarkers for post-mortem interval estimation: A comprehensive review of proteomic approaches in forensic science. J Proteomics. 2025 Jan 6;310:105326.

68. Yssouf A, Almeras L, Raoult D, Parola P. Emerging Tools for Identification of Arthropod Vectors. Future Microbiol [Internet]. 2016 Apr [cited 2025 Jul 29];11(4):549–66. Available from: https://www.tandfonline.com/doi/full/10.2217/fmb.16.5

69. Fang Q, Keirans JE, Mixson T. The Use of the Nuclear Protein-Encoding Gene, RNA Polymerase II, for Tick Molecular Systematics. Exp Appl Acarol [Internet]. 2002 May [cited 2025 Jul 29];28(1–4):69–75. Available from: https://link.springer.com/10.1023/A:1025389914156

70. Moreau CS, Wray BD, Czekanski-Moir JE, Rubin BER. DNA preservation: a test of commonly used preservatives for insects. Invertebr Syst [Internet]. 2013 [cited 2025 Jul 29];27(1):81. Available from: http://www.publish.csiro.au/?paper=IS12067

71. Krehenwinkel H, Fong M, Kennedy S, Huang EG, Noriyuki S, Cayetano L, et al. The effect of DNA degradation bias in passive sampling devices on metabarcoding studies of arthropod communities and their associated microbiota. PloS One. 2018;13(1):e0189188.

72. Stojan I, Trumbić Ž, Lepen Pleić I, Šantić D. Evaluation of DNA extraction methods and direct PCR in metabarcoding of mock and marine bacterial communities. Front Microbiol. 2023;14:1151907.

73. Lienhard A, Schäffer S. Extracting the invisible: obtaining high quality DNA is a challenging task in small arthropods. PeerJ. 2019;7:e6753.

74. Raupach MJ, Astrin JJ, Hannig K, Peters MK, Stoeckle MY, Wägele JW. Molecular species identification of Central European ground beetles (Coleoptera: Carabidae) using nuclear rDNA expansion segments and DNA barcodes. Front Zool. 2010 Sep 13;7:26.

75. Virgilio M, Backeljau T, Nevado B, De Meyer M. Comparative performances of DNA barcoding across insect orders. BMC Bioinformatics. 2010 Apr 27;11:206.

76. Nilsson RH, Ryberg M, Kristiansson E, Abarenkov K, Larsson KH, Kõljalg U. Taxonomic Reliability of DNA Sequences in Public Sequence Databases: A Fungal Perspective. Fairhead C, editor. PLoS ONE [Internet]. 2006 Dec 20 [cited 2025 Jul 29];1(1):e59. Available from: https://dx.plos.org/10.1371/journal.pone.0000059

77. Floyd R, Lima J, deWaard J, Humble L, Hanner R. Common goals: policy implications of DNA barcoding as a protocol for identification of arthropod pests. Biol Invasions [Internet]. 2010 Sep [cited 2025 Jul 29];12(9):2947–54. Available from: http://link.springer.com/10.1007/s10530-010-9709-8

